# Pervasive low-frequency vocal modulation during territorial contests in Eurasian Scops Owls (*Otus scops*)

**DOI:** 10.1101/2020.12.07.415117

**Authors:** Fabrizio Grieco

## Abstract

In several animal species, including birds, individuals are known to produce low-frequency vocalizations during aggressive interactions with conspecifics. In this study, I investigated territorial interactions between male Eurasian Scops Owls that occupy territories in a densely-packed area. The single-note hoot of the Scops Owl is generally assumed to be highly repeatable, however extensive recording of male-male interactions identified previously unrecognized variation in the hoots’ structure. Male Scops Owls gave hoots at a frequency lower than usual when engaging in short-distance contests with neighbouring males. Within-subject analysis revealed that the caller’s average hoot frequency was positively correlated with the distance from its rival. During contests, males gradually reduced their hoot frequency as they approached one another, perhaps reflecting changes in the degree of escalation. Male Scops Owls reduced their hoot frequency immediately after the rival initiated countersinging, and returned to their usual frequency range at the end of the contest. These findings demonstrate that vocal modulation is pervasive in social contexts in the Scops Owl, and suggests that individuals have full voluntary control of the structure of their vocalizations. This study confirms in part the findings of other authors’ experimental work, where male owls adjusted their vocal frequency when challenged by an opponent. However, that study suggested that vocal frequency would encode information about the caller’s body weight. In contrast, the present study revealed context-dependent fluctuations in hoot frequency, which may suggest that the hoot of the Scops Owl dynamically reflects the current motivational state of the caller during the contest.

## INTRODUCTION

Contest behavior in animals involves some of the most dramatic interactions and displays that we can observe. Aggressive displays are thought to have evolved as a substitute of fighting that enables individuals to assess the opponents’ fighting abilities, and settle disputes for resources without incurring the risk of costly injuries (Tinbergen, 1951; Maynard Smith & Price, 1973; Bradbury & Vehrencamp, 1998). Animals show marked variation in the type of threat signals both across and within species, particularly vocalizations (Searcy & Beecher, 2009; Bradbury & Vehrencamp, 1998; Morton, 2017). A common finding of research work in communication in animal contests is that individuals produce low-frequency sounds when engaging in disputes with conspecifics (Hill & Lein, 1987; Price, *et al*., 2006; Benedict, *et al*., 2012; Geberzahn, *et al*., 2009; Geberzahn & Aubin, 2014); for an overview see (Morton, 2017). There is a fundamental reason why vocal frequency plays such an important role in aggressive contexts, and that is the isometric scaling of the sound-producing mechanisms to body size. Especially in animals with indeterminate growth, and whose individuals vary significantly in size, like amphibians, larger individuals have a larger vocal apparatus, which in turn dictates the range in frequencies that those individuals can produce. Larger animals, with greater fighting abilities, could advertise their size or condition by giving vocalizations at a lower frequency that other, smaller conspecifics could not produce. The ancient relationship between call frequency, body size and aggression is thought to be conserved in birds and mammals, where different vocalizations are no longer an index of the body size but represent differences in aggressive motivation (Morton, 1977). In birds, the frequency of vocalizations correlates with body size or condition, both between species (Ryan & Brenovitz, 1985; Seddon, 2005; Bertelli & Tubaro, 2002; Price, *et al*., 2006; Cornec, *et al*., 2015; Wallschläger, 1980; Tubaro & Mahler, 1998) and within species (Appleby & Redpath, 1997; Brunton, *et al*., 2010; Mager, *et al*., 2007; Hardouin, *et al*., 2007; Hall, *et al*., 2013). Other studies, however, found no such correlation (Logue, *et al*., 2007; Cardoso, *et al*., 2008). Although the size dependence of frequency is probably more complicated by several physiological factors (Goller & Riede, 2013), it has strong theoretical (Fletcher, 2004) and empirical (Bowling, et al., 2017) support and suggests that animals could use frequency as a cue to body size or some other inherent quality of conspecifics in male-male competition contexts (Morton, 2017).

The monotonous, repeated call (henceforth “hoot”) of the Eurasian Scops Owl (*Otus scops*) is thought to serve two functions: to attract mates and to warn off rivals (Koenig, 1973). Some acoustic properties of Scops Owls’ vocalizations, like their rhythm and dominant frequency, are highly repeatable within individuals, even when measured across years (Koenig, 1973; Galeotti & Sacchi, 2001; Dragonetti, 2007; Muraoka, *et al*., 2009). Male Scops Owls give a whistle-like note during bouts that may last several minutes or even hours, which may lead to think that their “intended signal” is stable. However, Hardouin *et al*. (2007) showed in an experimental work that male Scops Owls adjusted their hoot frequency as a response to a simulated territorial challenge. The authors found that in natural populations hoot frequency was negatively correlated with body weight – the heavier the individual, the lower its vocal pitch. In their experiment, male owls were stimulated with hoots mimicking light and heavy individuals. The authors found an association between the weight of the focal subjects and the degree of response in terms of frequency change. When the playbacks simulated heavier intruders, heavier males tended to give lower-frequency hoots than when they when the playbacks simulated lighter individuals. The relationship was inverse for lighter males, that is, lower-frequency stimulation resulted in higher tones. The authors concluded that the hoots encode information about the caller’s body weight, interpreted as an index of body condition, and proposed that individuals produce low-frequency vocalizations to “sound” larger than they were when responding to a frequency indicative of heavier individuals. Further work showed that males with lower voice pitch have higher breeding performance (Hardouin, *et al*., 2009).

There are, however, questions that remain to be answered. First, there is still no evidence that frequency adjustments similar to those described by Hardouin *et al*. (2007) also occur in dyadic interactions of freely-interacting Scops Owls. Second, Hardouin *et al* (2007) measured the first ten hoots of the focal males after playback stimulation. This corresponds approximately to a 30-second interval; therefore, it is not known whether the response to a territorial challenge was consistent over a longer time. Playback experiments are often carried out with limited numbers and duration of stimuli (Hopp & Morton, 1998) and may provide a partial view of the natural variation in the response. In general, time-related changes in acoustic properties of vocalizations during territorial contests are poorly documented and this makes it difficult to assess their repeatability and reliability as signals (Biro & Stamps, 2015). Besides, Hardouin and co-workers (2007) viewed their findings as preliminary, calling for further research, yet no other study so far appears to have provided new information. Scops Owls provide an excellent opportunity to investigate their communication system because they are highly responsive to the voice of conspecifics (Galeotti, *et al*., 1997) and sometimes form aggregations (Grieco, 2018) which makes it possible to observe territorial disputes between known individuals. At high densities, territory-holding males are more likely to come into contact and engage in disputes. In this study, I explored the dynamics of spontaneous countersinging interactions between Scops Owls and formulated the following three predictions:

### Prediction 1: Occurrence of low-frequency vocalizations

Low-frequency vocalizations are more likely to occur when two opponents approach one another, based on the assumption that a shorter distance between opponents reflects a higher degree of escalation. For this reason, a positive correlation should appear between the hoot frequency of the focal individual and the distance from the nearest neighbour (henceforth the *Rival*).

### Prediction 2: Frequency decrease after the rival’s initial countersinging

If individuals monitor the calling activity of other males in the neighbourhood, it should be possible to detect the start of a low-frequency contest. Therefore, males that were singing were predicted to switch to a lower frequency when a nearby male initiated countersinging.

### Prediction 3: Frequency increase at the end of an interaction

A territorial interaction was assumed to end when one of the two opponents left, including when that individual resumed calling further away from its original position. Hoot frequency was predicted to increase after that moment.

## METHODS

### Study area

The study was carried out in the State Nature Reserve “Metaponto”, located along the Ionian coastline in the administrative region of Basilicata, Italy. The main habitat is a pine (*Pinus Alepensis*) plantation, alternated with sparse human settlements, mainly beach resorts and camping areas. Scops Owls cluster in groups and leave large portions of apparently suitable habitat deserted. The study population occupies an area of approximately 8 hectares, the larger part being within a camping area. Male owls hold relatively small, contiguous territories, with a mean distance between territory centres being under 50 m, and calling sites of different males often overlapping (further details in Grieco (2018)).

### Vocalization recording

Scops Owl hoots were recorded in calm and dry nights between 00:00 and 04:30 in July and early August in the years 2017 and 2018. That period of the year corresponds to the time that the pairs raise their broods (besides the breeding pairs, some resident males are thought not to breed every year). Vocalizations were recorded using a Telinga Pro 8 MK2 stereo microphone mounted on a parabolic dish with a diameter of 52 cm, and a Marantz PMD 661 MK2 digital sound recorder. Recordings were stored in WAV files at a sample rate of 44.1 kHz, with 16-bit depth. Often a recording session (henceforth *Recording*) started when one or more owls were already singing, and in any case, stopped when the focal individuals were silent for one minute. An effort was made to record as many individuals and as many territorial interactions as possible on each night. Because the study intended to focus on within-individual variation in calling behaviour, recording multiple, repeated measures of the same individual was given higher priority over recording more individuals in a larger area. The position of individuals was noted on a map, together with changes in their position, observation of flights and agonistic interactions that occurred during the recording session. Throughout the text, I assume that the males’ calling activity was mainly related to male-male competition rather than mate attraction. This assumption is supported by the observation that males often stop calling shortly after their nearest neighbours do so (unpubl. data).

### Individual identification

The hoots of Scops Owls are highly repeatable within individuals (Koenig, 1973), even across years, and one can use their acoustic properties to identify individuals (Galeotti & Sacchi, 2001; Denac & Trilar, 2006; Dragonetti, 2007). This work focuses on male-male interactions, as females were mostly silent at that time of the year. Males were identified using two diagnostic features: the dominant frequency measured at peak amplitude (see below for details) and the time between the onset of consecutive hoots. In total, I analysed 343 recordings involving 19 identified males, singing either solitarily (80 recordings with one identified male, 23%) or in groups of two up to five individuals (263 recordings, 77%). Males identified in at least five recording sessions and for which their distance from a neighbour was known were selected as focal males *(n* = 12). Other identified males *(n* = 7) occupied their territories at the margin of the study area and were underrepresented in the recordings.

### Frequency measurements

Hoot frequency was analysed with Audacity 2.0. For each hoot, I extracted the frequency at maximum amplitude (henceforth *Peak frequency*). The assumption here is that peak frequency reliably measures the main component of the signal that the caller intended to send to its audience. When the power spectrum showed two peaks that differed by less than 1 dB in amplitude, the average frequency was taken (as a side note, here the amplitude is not meant as sound pressure level; instead, it is given relative to the full-scale reading, where 0 dB is the maximum signal that can be recorded by the device without frequency clipping). In all cases, the following spectrogram parameters were used: FFT window length 4098 samples, Hanning window, frequency resolution 10.8 Hz, temporal resolution 0.093 s.

### Prediction 1

To test whether males reduce their hoot frequency when in the proximity of other males, I considered the average peak frequency calculated for each recording and individual male. The Nearest Neighbour Distance (NND) was calculated as the minimal distance to any male that called simultaneously during that recording. Because the average hoot frequency and NND were not normally distributed, the data were transformed; Peak frequency was square-root transformed, and NND was log-transformed. For each focal individual, I then computed the regression coefficients *B* of the transformed frequency on the transformed NND. For the individual males that were observed in both study years, the average coefficient was entered in the analysis. The regression coefficients were tested with a one-sample t-test against the average of zero that was expected based on the null hypothesis of no relationship between the two variables.

### Illustrative dataset

To illustrate the extent of the changes in the owls’ call structure, I selected a number of recordings, henceforth called *Low-Frequency Contests* (LFCs), that fulfilled two criteria: (1) The average peak frequency of at least one of the individuals changed by at least 22 Hz for 30 or more seconds relative to the overall average for that individual; and (2) occurrence of events that suggested that two or more males were interacting, including the following: a male approaching another one, and/or initiating countersinging, noisy wing-clapping, rapid movements in trees, chases and physical confrontation. The 22 Hz threshold corresponded to two times the frequency resolution of the spectral analysis. Although it may sound arbitrary, the resulting selection did not leave out any episode of apparently aggressive interaction. In total, I selected 44 LFCs (20 in 2017 and 24 in 2018). Throughout this work, I used the term “contest” which, according to Briffa and Hardy (2013) is “a direct and discrete behavioural interaction that determines the ownership of an indivisible resource”, and does not assume that the interaction escalates to physical contact.

### Prediction 2

To test whether male owls react to other males as soon as those males initiate their calling activity, I considered a group of episodes where the nearest neighbour initiated calling while the focal male had been calling for at least one minute without interruption. For this data set *(n* = 7 males, 21 episodes), the prediction was that the focal males’ hoot frequency would decrease after the rival’s initial hoot. For each episode, I extracted twenty hoots of the focal male: the last ten hoots before the initial hoot of the rival (henceforth called Hoot 1 to Hoot 10), and the first ten calls after the initial hoot of the rival (Hoot 11 to Hoot 20). Hoot frequency was normalized by subtracting the 20-point average from each value. Change Point Analyzer (Taylor Enterprises, 2003) was used to detect the change point in the 20-point time series. The change point was expected to occur at Hoot 11 of the focal individual. The software combines Cumulative sum charts (CUSUM) and bootstrap methods to simulate a large number of bootstrap samples based on the data. The bootstrap samples represent random reorderings of the data that mimic the behavior of the CUSUM if no change has occurred. The software outputs the detected change points and their confidence intervals. The following parameters were used: CUSUM, 10000 bootstrap samples, 90% Confidence Levels for inclusion of change points in the results, 95% Confidence Intervals around the change point. The analysis was run for each individual male separately; if more episodes were available per individual, the averages were computed for each hoot number (1 to 20). The significance level was corrected using Bonferroni’s alpha (with *n* = 7 males, α = 0.007).

### Prediction 3

To test whether Scops Owls call at higher pitch after an interaction ends, fragments of interactions were considered when either the focal male or its rival left the current site and resumed calling at a greater distance within five minutes. Hoot peak frequency was contrasted in the following 30-seconds intervals: *1 – Baseline*: from 120 s to 90 s before one of the two rivals left; *2 - Pre-Leaving*: a 30-s interval immediately before one of the two individuals left; *3 - Post-Resuming*: from 30 s to 60 s after the male that had left resumed calling within five minutes. To test for between-interval frequency changes, I performed a Friedman repeated-measures test and Nemenyi post-hoc tests using the Real Statistics Resource Pack software (Zaiontz, 2019). All tests were two-tailed and, if not otherwise stated, the significance level was set to 0.05.

## RESULTS

Scops Owls at Metaponto often sing together with their conspecifics. On average, only 10.1% ± 7.3 (SD) of the recordings in which a male was singing were in fact cases of solitary singing *(n* =12 males). Fifty-nine percent of the recordings included two identified males singing simultaneously, while 13% included three males and 4% four or five males. Note that those figures underestimate the total number of males singing simultaneously because they do not include unidentified males or males that sang outside the study area. Overall, the average distance between a singing male and its nearest neighbour was 34.2 ± 7.2 m *(n* = 12) and ranged from a minimum of 1.8 m to a maximum of 181 m.

### Prediction 1: Occurrence of low-frequency vocalizations

Male Scops Owls called at a lower frequency when they were closer to their rivals. Figure 1A shows the individual’s hoot frequency expressed as the deviation from the overall average, plotted against the distance of the focal individual from the nearest neighbour. The effect of distance from the neighbour was negligible at more than 20 m. When a male was at less than 10 m from a neighbour, its hoot frequency dropped by 30.7 ± 18.2 (SD) Hz *(n* = 12) compared with the individual’s average frequency. The within-focal individual analysis revealed that hoot frequency was positively associated with distance from the nearest neighbour NND (regression coefficients of Average hoot frequency on NND: +2.30 ± 1.73 (SD) *(n* = 12 owls) and the coefficients were significantly greater than zero (one-sample t-test, t = 4.60, df = 11, *P* = 0.0008; 95% Confidence limits 1.20 – 3.40; effect size Cohen’s *d* 1.33). The relationship was not the result of data points of specific pairs of interacting males clustering in specific regions of the distance-frequency space (Figure 1B). Furthermore, when considering the few individuals that were observed in the two years of the study, the distance-frequency relationship was consistent between years (see an example in Supplementary Figure S1). It could be argued that the observed positive association would appear if subsequent recordings were correlated, for example when two recordings occurred at few minutes between one another. However, average hoot frequency and NND were not autocorrelated when considering each focal individual (Autocorrelation functions, lags 1 to 5, always *P* > 0.05).

**Figure 1.**
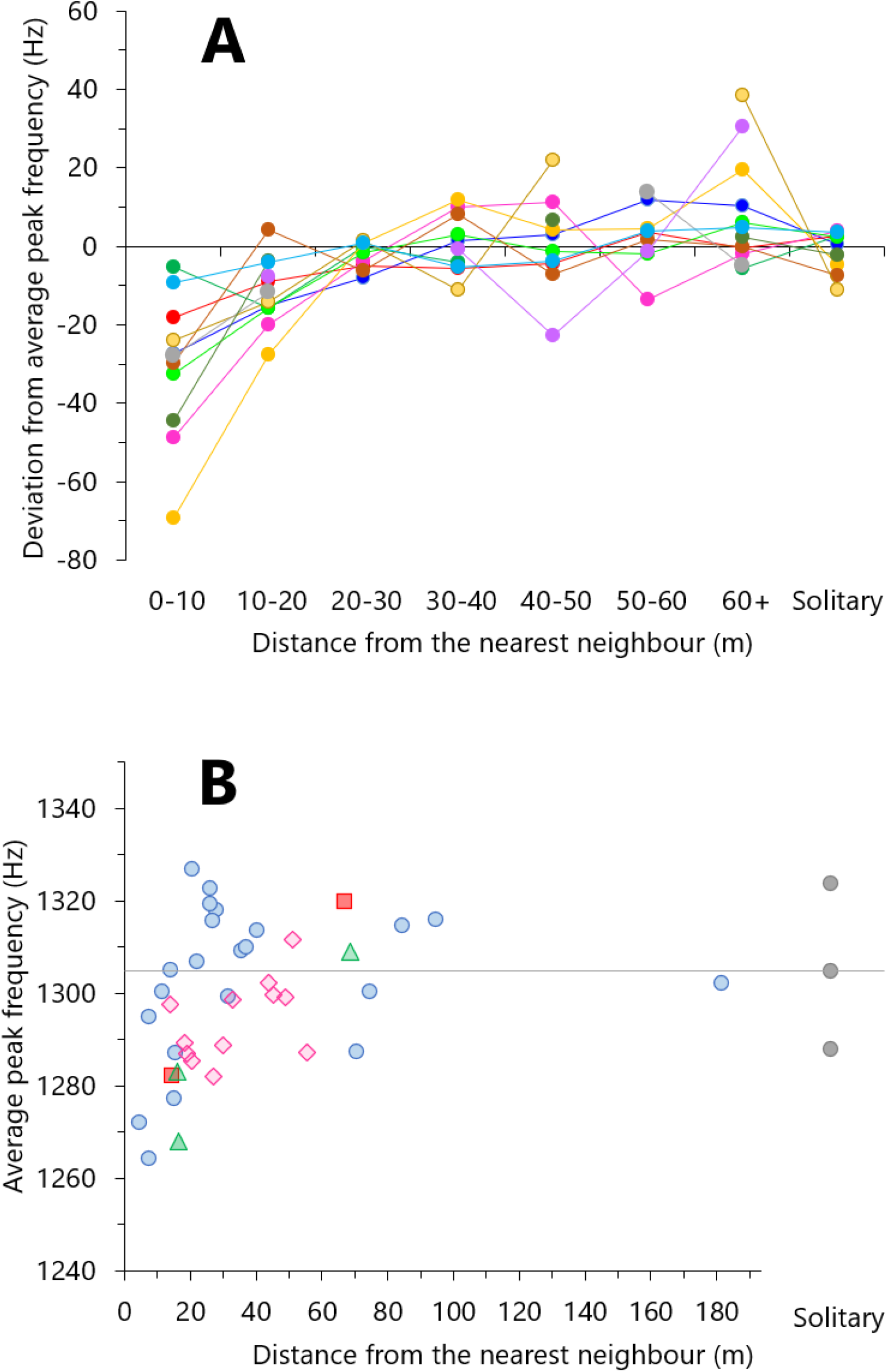
The relationship between the average peak frequency of male Scops Owl hoots and the distance from the nearest neighbour. **A.** Each data point represents the average for each category of distance, for 12 focal males (indicated with colours). Frequency is expressed as the deviation from the individual’s overall average, that is, negative values indicate hoot frequency lower than the average. **B.** Peak hoot frequency of male *NorthGreen* in 2018 plotted against the distance from its nearest neighbour. Each data point represents one recording session, symbols and colours indicate unique focal-rival male pairs. The frequency is the per-session average, and the distance is the minimal distance measured in that session. Dots at the right side indicate the average hoot frequency during solitary singing. The horizontal line represents the average hoot frequency based on all recordings.

The charts in Figure 1 also show the male’s hoot frequency measured when they called solitarily. Hoot frequency was significantly higher during solitary singing than when the male was at less than 20 m from its rival (deviation from overall average: solitary singing, −0.9 ± 5.2 Hz; At 20 m or less from a rival: −19.1 ± 11.4 Hz; Wilcoxon paired test, W = 0, W_crit_ = 8, *P* < 0.05; Figure 1A). Therefore, it is concluded that prediction 1 was met. When Scops Owls engaged in territorial interactions at a short distance from one another, their produced vocalizations at a frequency lower than usual.

### The course of a short-distance interaction

So far, we have examined averages collected in single recording sessions, which may last up to several minutes. What happens during a single session? I give here a brief account of a low-frequency contest between owl *Blue*, a resident breeding male, and owl *Green*, a resident, non-breeding male, recorded on July 22nd, 2017. Note that these two birds had similar vocal frequencies (averages 1995 Hz and 1088 Hz, respectively). In this episode, depicted in Figure 2, *Blue* initiated calling loudly at its typical broadcasting frequency (Figure 2A, B). Two minutes later, Green started calling within ten meters from Blue (Figure 1C). From that moment, *Blue* called at a progressively lower frequency (Figure 2A, C). At four minutes, Blue approached Green; this event apparently triggered an even stronger reduction in hoot frequency in both males. At 7’45’’ the birds made short flights and jumps through the foliage, apparently chasing each other; possibly physical confrontation occurred given the noise of wing flapping in the tangle of branches. Hoot intensity decreased to such an extent that the males could only be heard when standing a few meters away (Figure 2D). The interaction went on for several minutes and eventually the two males left; first *Blue*, then *Green*, apparently ending the interaction. Overall, *Blue* exhibited a stronger reduction in frequency than *Green* and this was consistent across contests. Throughout the study period, I observed six LFCs between those two males, and in all six episodes, *Blue* reduced frequency on average 60 Hz more than its rival (range 20 - 81). Figure 2E gives an overview of frequency changes during LFCs. On average, hoot frequency dropped to a minimum of 69.4 ± 40.6 (SD) Hz below the individual’s average (range: 24 - 161 Hz, *n* = 12). Additional examples of time-related changes in hoot frequency are shown in Supplementary Figure S2.

**Figure 2.**
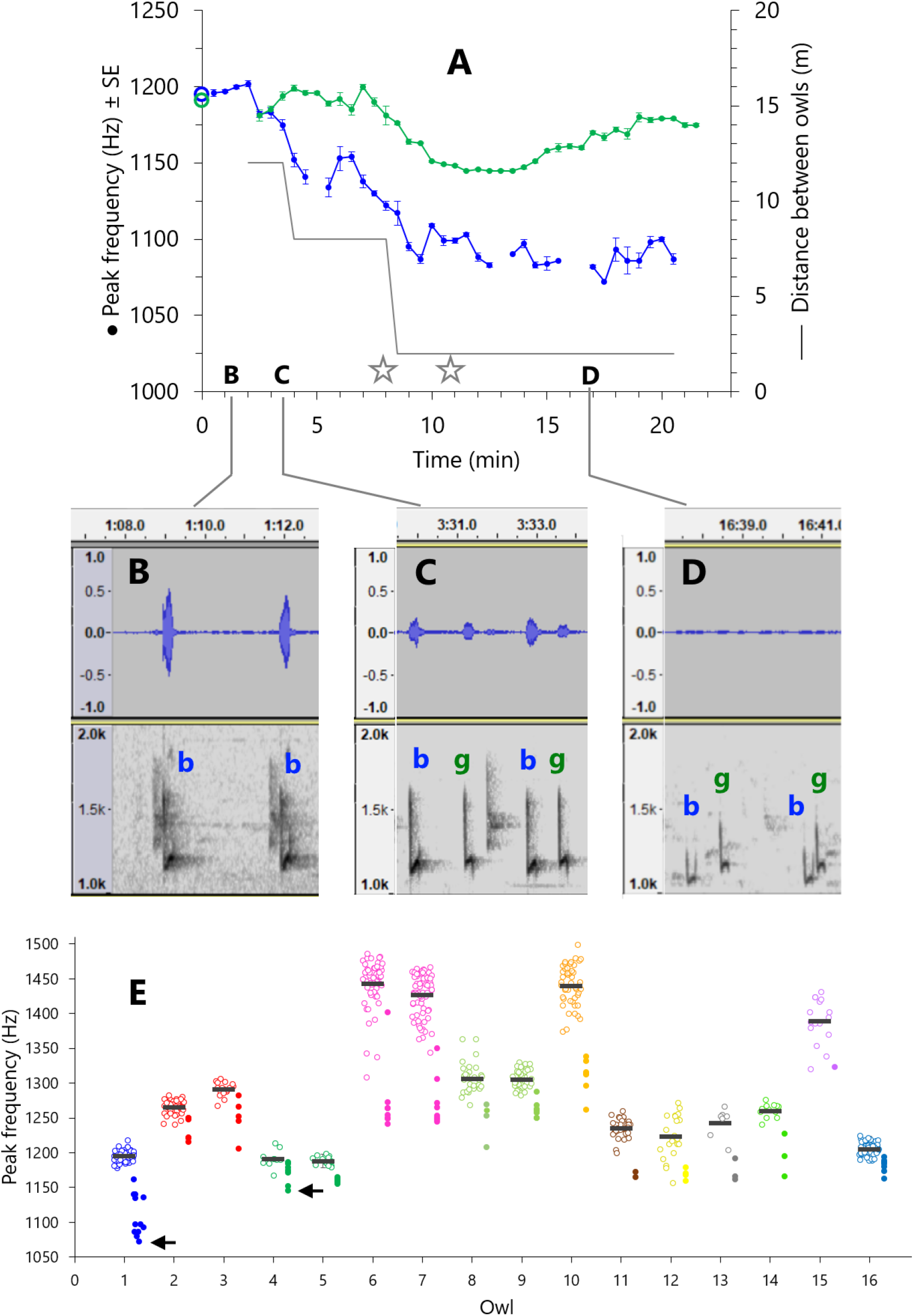
**A.** Temporal variation of hoot frequency of two male Scops Owls, *Blue* and *Green*, recorded on July 21, 2017. Peak frequency is averaged over 30-s intervals (see Methods). The grey line indicates the distance between the two males. The stars indicate when chases were observed. The circles on the y-axis indicate the overall average hoot frequency. **B-D**: Representative oscillograms (top) and spectrograms (bottom) of hoots at three time points. Lowercase letters indicate the calls of *Blue* (b) and *Green* (g). In **C**, a third male, *Fuchsia*, called on the background. Note the marked decrease in call amplitude and the change in the shape of the spectrograms in **C** and **D**. The asymmetry in frequency change between the two opponents was observed in six contests (see text). **E**: Overview of frequency changes during LFCs for each individual male and year. Different colours represent individual males, recorded in one year (e.g. Owl 1) or two years (e.g. Owl 2-3). Dots represent the lowest frequency measured in 30-second intervals during LFCs (see Methods). Two small arrows indicate the values for males *Blue* and *Green* during the contest pictured in **A**. Note that not all dots lie well below the average frequency; they refer to contests when the male did not reduce much its frequency, while the opponent did so. Circles indicate the average hoot frequency in the remaining (non-LFC) recordings, the black segments indicate the grand averages.

### Prediction 2: Frequency drop after the rival’s initial countersinging

Scops Owls switched to lower-frequency vocalizations immediately after the nearest neighbour initiated countersinging. Figure 3A shows the first response of owl *Blue* to the first hoots of owl *Green* (see also Figure 2A). After *Green*’s first hoot, Blue’s frequency dropped by approximately 20 Hz and remained stable in the new range 1175 to 1190 Hz. I then analysed similar episodes for each focal male, using the 20-hoot sequences around the first hoot of the rival male (see the methods; Figure 3B). For five of the seven focal males, bootstrap-based change point analysis detected the point of frequency change at Hoot 11 (95% Confidence Interval: Hoot 10 to 11 for four males, 11 to 14 for one male). For the remaining two males, the change point was not significant after Bonferroni correction. When all episodes were lumped together *(n* = 7 males, 21 episodes), the change point was significant at Hoot 11 (*P*< 0.01, 95% Confidence Interval: 9, 11). The largest drop in frequency was found between Hoot 10 and Hoot 11 (−43.3 ± 49.1 Hz (*n* = 7); Wilcoxon paired test, contrast Hoot 10 vs Hoot 11: *W* = 0, *W*_crit_ = 2, *P*< 0.05; Figure 3B). Therefore, prediction 2 was met. It was demonstrated that male Scops Owls reacted to another male’s initial countersinging by reducing their vocal frequency immediately after hearing the rival’s first hoot.

**Figure 3.**
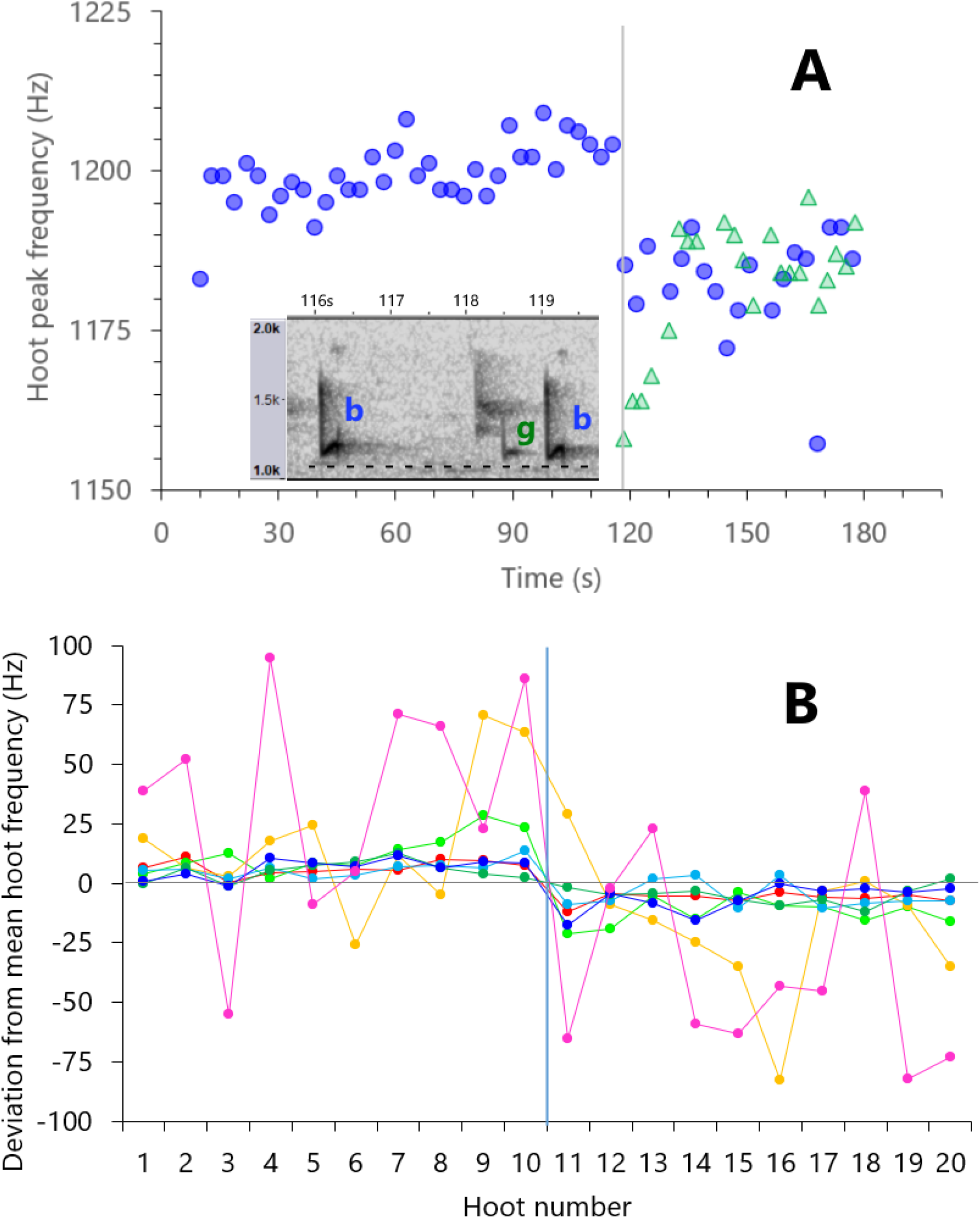
Response to initial countersinging. **A.** The peak frequency of single hoots of owl *Blue* (circles) measured at around the time that its main opponent, *Green*, started calling (triangles). This is part of the episode depicted in Figure 1A, zoomed in at about two minutes from the start. The inset shows the spectrogram of the hoots of Blue (b) immediately before and after the first hoot of Green (g). The horizontal dotted line helps compare the frequency in the spectrograms. Owl *Blue* reduces its hoot frequency within a fraction of a second from the first hoot of *Green*. Note that the first hoot of *Blue* and first few hoots of owl *Green* have a low frequency; that is typical in Scops Owls’ hoot trains. **B.** Hoot frequency changes in seven males around the time that another male started singing. The plot shows the change in the frequency of ten hoots of the focal male immediately before the opponent’s first hoot (1-10) and the focal’s next ten hoots (11-20). Frequency is normalized by taking the difference from the 20-hoot average. Data points represent averages calculated per individual. The frequency change between Hoot 1-10 and Hoot 11-20 was −26.7 ± 23.0 (SD) Hz.

### Prediction 3: End of the escalated contest

When one of the contestants leaves the site of the interaction, the level of arousal in both individuals is expected to decrease, and vocal frequency should return to typical levels. Figure 4A pictures one of such episodes. Two males, *NorthGreen* and *Red*, engaged in a low-frequency contest for about twenty minutes. Eventually, *NorthGreen* left and resumed singing at about 80 m from *Red*. Figure 4A shows the fast increase in the hoot frequency of *NorthGreen* and, to a lesser extent, of *Red* who went on calling. When taking all episodes together, it turns out that one of the two, or both, called at significantly higher frequency after the end of a short-distance interaction (Figure 4B, timepoints 2 vs. 3). The sample size *(n* = 7 males, 13 episodes) does not allow to test the prediction separately for only those males that left, or only those that stayed. The data, however, provide support to the view that male Scops Owls can quickly resume calling at their regular vocal frequency at the end of an escalated contest; such prompt recovery also suggests that graded reduction in frequency was not due to exhaustion.

**Figure 4.**
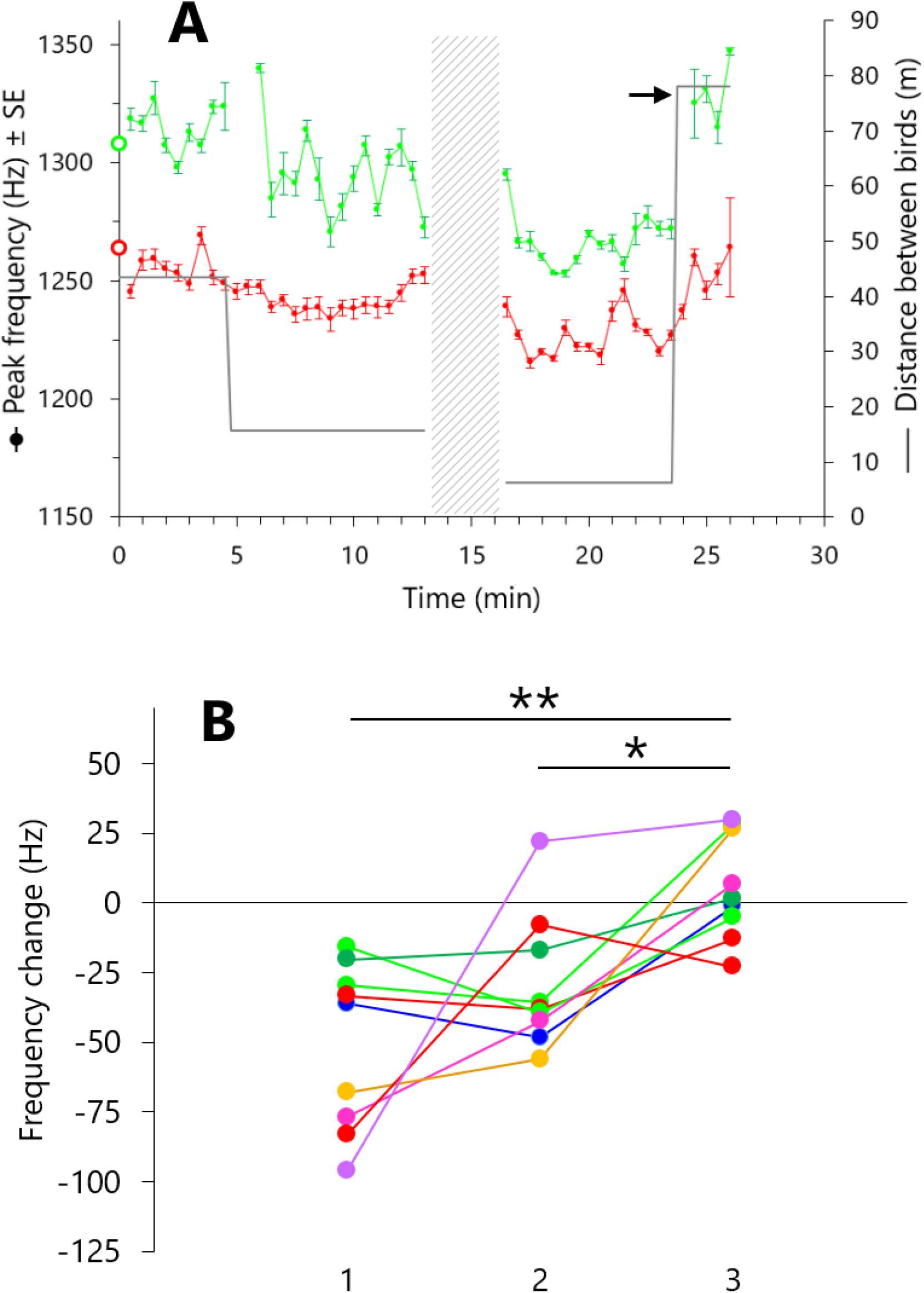
**A.** Temporal variation in hoot frequency of two male Scops Owls, *NorthGreen* and *Red*, on July 17, 2017. Data are shown as in Figure 2A. The arrow indicates when NorthGreen left and resumed calling further away. The shaded area indicates the time when the calls were not recorded because other owls were recorded elsewhere. Note the drops in hoot frequency as the two opponents get closer to each other. At about 24 minutes, hoot frequency quickly returned to regular levels, particularly for owl NorthGreen, after the interaction ended. **B.**Variation in call peak frequency of male Scops Owls at three intervals around the moment that the focal male (or its current rival) left and resumed calling elsewhere. The frequency change is the deviation from the average frequency of an individual in a particular year (see Methods and Figure 1E). **1**: 120 s to 90 s before one of the two rivals left. **2**: 30 s up to when one of the two rivals left. **3**: 30 s to 60 s after the leaving bird resumed calling (max 5 minutes after leaving). Colours represent individual males. Two males (indicated with red and light green dots) were observed in both study years. Males called at higher frequency after the end of the interaction (Friedman repeated measures test: Q = 11.14, df = 2, P= 0.004. Nemenyi post-hoc tests: **1** vs **3**, Q = 4.53, *P* = 0.004; **2** vs. **3**, Q = 3.40, *P* = 0.04; **1** vs. **2**, Q = 1.13, n.s. Tests were performed using per-individual averages; n = 7 owls). From time point **2** to **3**, the distance between the two rivals increased from 15.5 ± 11.9 (SD) m to 64.6 ± 19.4 m.

## DISCUSSION

The most significant findings can be summarized as follows:

1. Extensive recording of spontaneous dyadic territorial contests revealed a large amount of variation in the call structure in a bird species that is generally assumed to produce highly repeatable vocalizations. As a neighbour was progressively closer to the focal individual, and thus as the level of threat increased, males gradually produced low-frequency hoots.
2. Males appeared not to signal at the maximal level at the beginning of an encounter, rather they showed varying degrees of escalation and de-escalation during the course of the interaction. However, sudden drops in hoot frequency occurred, apparently associated with the initial countersinging by the opponent, and also with chases and aggressive displays that were observed during the contest. Scops Owls returned to their typical vocal frequency when the contest ended. These observations suggest that Scops Owls have voluntary control of their vocalization frequency.

### Hoot frequency as a graded signal?

Understanding communication requires us to examine complex, varying signals (Patricelli & Hebets, 2016) and not to ignore time-related changes in behaviour (Biro & Stamps, 2015). Although the significance of low-frequency vocalizations in competitive contexts has been established in a number of species (see Introduction), to the best of my knowledge I am not aware of time-related changes in vocal frequency in the course of a contest. The fluctuations in hoot frequency like those in Figure 2A and 4A are something of a puzzle; they resemble one-to-one dialogues rather than typical signal broadcasting to a broader audience and suggest that something profoundly changes during the course of the interaction. The experimental work by Hardouin *et al*. (2007) suggests that an individual challenged by a rival communicates information about its own body weight (or any unknown variable correlated with it). If vocal frequency were a proxy for individual quality only, and assuming that quality does not change over a short time, then one would expect that signal to be rather stable during a contest, all else being equal. This seems in contrast with the high variability in hoot frequency both between different contests and within contests found in this study. Perhaps the patterns like that in Figure 2A resulted from males attempting to sound progressively heavier at increasing levels of escalation, but then how can such marked within-individual variability be effectively used as an indicator of relatively fixed qualities? Patel *et al*. (2010) found that in male black swans (*Cygnus atratus*) vocalization frequency was correlated with body size across individuals. However, they also found that males had high within-individual variability in vocalization frequency, and concluded that the reliability of vocalization frequency as a signal of body size was low. Similarly, Wagner (1992) found that experimentally-induced reduction in call frequency of cricket frogs made call frequency a less reliable signal of body size.

Observed differences in the behaviour between opponents suggest that the typical frequency of an individual does not predict the frequency change during a contest. Two individuals, *Blue* and *Green*, with virtually the same vocal frequency, and therefore expected to be of equal weight based on what found by Hardouin *et al*. (2007), showed a consistent asymmetry in the frequency change across several contests (see Fig 2A’s caption); therefore, the correlation with their original frequency was lost. Owl *Blue*, a nest owner, reduced hoot frequency to a greater extent than its main rival *Green*, a non-breeding resident, and *Red*, another non-breeding neighbour. Owl *Blue* called generally louder than Green, and at the end of interactions he often returned to the centre of its own territory and resumed calling at full intensity. In contrast, owl *Green* never did so, as would disappear after those encounters. This might indicate that *Blue* was somewhat dominant in the area. Could a consistent asymmetry in hoot frequency have something to do with betwee-individual variation in breeding status or social dominance? Clearly, an appropriate sample of dyads involving individuals with similar vocal frequencies, and preferably of known body weight, is needed in order to have a clear answer. However, those observations suggest that changes in vocal frequency might also be associated with the context, and not only with a relatively fixed quality of individuals.

When a signal shows significant temporal changes, there are at least two factors that may help to explain variation: physiological condition (i.e. energetic state) and motivation, that is, the willingness of an individual to persist in a conflict. The former seems unlikely to be important here. The fast changes in vocal frequency, particularly the increase observed at the end of the contests, suggest that prolonged calling activity did not significantly reduce the energy reserves of the opponents. Therefore, vocal frequency probably provides little information on anything associated with the caller’s immediate energetic state. Graded signals have been shown to indicate varying levels of motivation (Bradbury & Vehrencamp, 1998; Vehrencamp, *et al*., 2014). If motivation results in winning more contests, individuals could be selected to send, and attend to, signals of aggressive motivation (Hurd & Enquist, 2001). I speculate that Scops Owls might signal their current motivation to enter into and persist with contests by producing low-frequency vocalizations. This hypothesis would hold despite the correlation found between body weight estimate and vocal response to intruders (Hardouin *et al*., 2007) if heavier individuals also had a higher aggressive motivation. For example, when heavier individuals occupy richer territories and this influences how they value their resources. In that case it would be difficult to disentangle the influence of body weight from that of motivation.

### Issues and open questions

An intriguing fact that deserves to be investigated is that male Scops Owls sometimes reduced hoot intensity, especially when engaging in long contests at close proximity (Figure 2A). This is contrary to what one would expect to occur if vocal performance increases with the level of escalation, and loud vocalizations and high vocal performance are important in deterring rivals and winning contests (Gil & Gahr, 2002; de Kort, *et al*., 2009). Low-amplitude vocalizations have been associated with aggressive motivation in a number of bird species, mostly songbirds (Dabelsteen, *et al*., 1998; Ballentine, *et al*., 2008) but also in other taxa (Ręk & Osiejuk, 2011; Reichard & Welklin, 2015); for a recent review see (Akçay, *et al*., 2015). There are several hypotheses that explain “soft calling” as adaptive behaviour (Dabelsteen, *et al*., 1998; Akçay, *et al*., 2015; Vargas-Castro, *et al*., 2017). However, in some situations, loud vocalizations might be difficult to produce because of physical or physiological constraints. Following a basic principle, animals cannot radiate high amplitude sounds with wavelengths much greater than those animals are (Bradbury & Vehrencamp, 2011). Small birds can produce sounds lower than 1 kHz, but their vocal tract acts as a high-pass filter which limits the lowest resonance frequency. Therefore, at low frequencies, a positive correlation between amplitude and frequency may appear (Ritschard & Brumm, 2011; Gaunt & Nowicki, 1998). It follows that birds may be able to lower their fundamental frequency only at the cost of reduced call intensity. This hypothesis is supported by the fact that a reduction in hoot amplitude co-occurs with a drop in frequency especially in low-frequency individuals, not high-frequency individuals (unpubl. data). Anyway, further investigation is needed to understand the relationship between intensity and frequency of vocalizations in this species.

In conclusion, the hoot of the Scops Owl seems to be a signal more complex than is currently recognized, and whose regulation and function are not entirely clear. Clearly, further research is needed that can illuminate questions about mechanistic and functional aspects of vocal communication in this species. I list here a few classes of questions: 1) *Inferring causality in the interaction*. The co-variation in frequency observed in several contests suggests the vocal behaviour of one individual is causally related to the subsequent change in behaviour of its opponent. The two individuals might attend to continuous variation in auditory information and assess each other’s states during the interaction and, on the basis of that information, respond with further escalations or de-escalations. 2) *Analysis of dyads*. Previous observations suggest that individuals respond more strongly to one neighbour than to another, and the same individual may show different behaviour in two years, perhaps as a result of variation in social status (e.g. breeding vs. non-breeding vs. floater). Understanding such asymmetries requires looking at low-frequency contests as properties of the pair of opponents (Logue & Krupp, 2016) and following individuals in multiple years. 3) *Analysis of contest duration and outcome*. A crucial question is what are the factors that affect the duration and the outcome of contests. Why do some contests last a few minutes while others up to one hour? So far, there is little evidence that males who call at lower frequency are more likely to make their opponent retreat (unpubl. data), however the sample currently available is too small to support any conclusion. 4) *Network approach*. Finally, Scops Owl aggregations are an ideal system for the study of communication in a network perspective. Although this work focused on dyadic interactions, multi-party contests also occur. This, combined with the observation that silent individuals attend to calling contests, raises the possibility that conspecifics could pay attention to and assess the relationship between the interacting individuals (McGregor, 2005).

## Acknowledgments

I wish to thank the staff of the Camping Park “Riva dei Greci” at Metaponto, especially Massimiliano Cospite for his interest in this study and permission to do fieldwork within the camping area.

## Data Availability Statement

The data that support the findings of this study are available from the corresponding author upon reasonable request.

## Supplementary material

**Supplementary Figure S1**. Peak hoot frequency of male *NorthGreen* plotted against distance from the nearest neighbour, in two years 2017 (open circles) and 2018 (dots). The horizontal line represents the per-year average hoot frequency. For simplicity, one line is drawn given that the average frequency was very similar between years (2017: 1306 Hz; 2018: 1305 Hz).

**Supplementary Figure S2**. Examples of the within-contest time-related variation in hoot frequency (averages ±SE). In both episodes, the largest frequency change coincided with the minimal distance between the rivals, indicated with the grey line. Open circles at the left indicate the individual’s overall average hoot frequency. **A**. Co-variation of hoot frequency in two contestants, owl *Fuchsia* (higher frequency) and *Orange*. The stars indicate chases or wing-clapping (see Methods). **B.** Hoot frequency recovery at the end of a short-distance interaction between owl *NorthGreen* and *Green*. The interaction took place at the territory boundary. At about nine minutes, *Green* flew back to the centre of its territory and resumed calling, while *NorthGreen* did not move and kept calling.

**Supplementary Audio File S1**. Fragment of audio with the calls of two male Scops Owls corresponding to Figure 3A where owl *Blue* reduced its call frequency after owl *Green* initiated countersinging. This occurs at approximately 19 sec.

**Supplementary Audio File S2**. Fragment of audio with the calls of two male Scops Owls, *Blue* and *Green* where both individuals call at low frequency. This corresponds to about 8 minutes (see the first star in Figure 2A). At the end of the file, noisy wing clapping indicates the beginning of a chase.

