## Supplementary figures and images for "Pervasive low-frequency vocal modulation during territorial contests in Eurasian Scops Owls (*Otus scops*)"

### Supplemental Figure 1

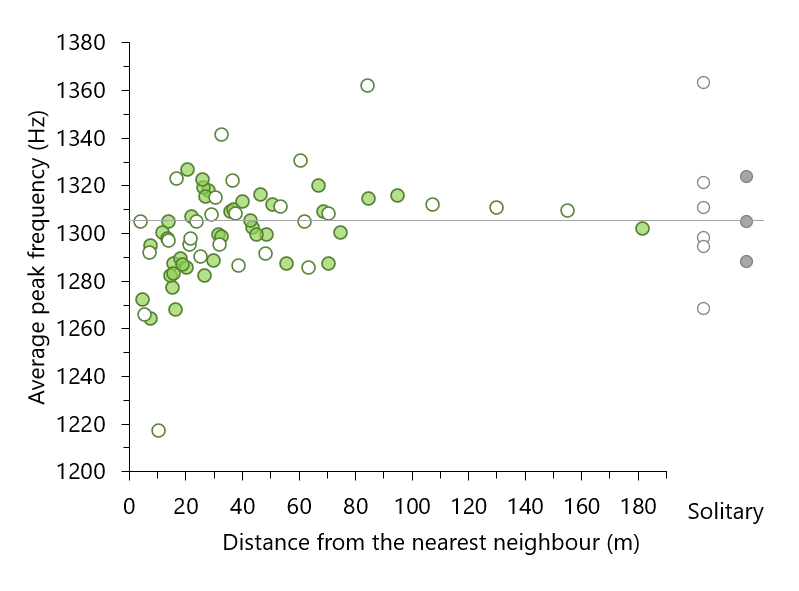

### Supplemental Figure 2

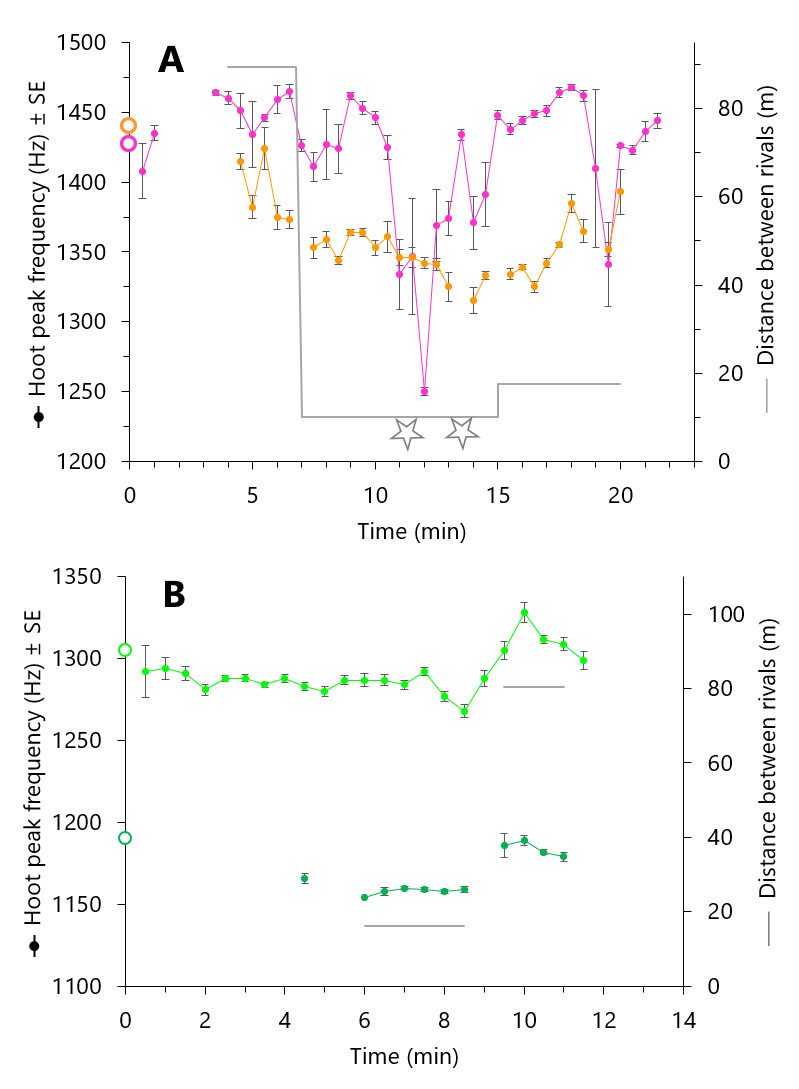
